# Genetic basis of glycine and L-serine toxicity in *Staphylococcus aureus* and the case for glycine as an antibiotic adjuvant

**DOI:** 10.64898/2026.06.08.730841

**Authors:** Tyler Brown, Jenna Barnum, Shalee Killpack, Joel Griffitts, Eric Wilson

## Abstract

The toxicity of the amino acids glycine and L-serine at high concentrations in bacteria was discovered decades ago. In this work, we used deep transposon insertion sequencing (Tn-seq) experiments to determine the genes necessary to tolerate excess L-serine, diglycine or glycine in the human pathogen *Staphylococcus aureus*. Our results indicate that intracellular accumulation of specific counterbalancing amino acids—such as alanine in excess glycine—is the primary mechanism of resistance to amino acid toxicity. Consistent with this model, specific amino acid and peptide uptake transporters were required for fitness in each treatment; the peptide transporter DtpT was crucial for fitness in excess L-serine or glycine, and the alanine transporter AapA was essential in diglycine. Tn-seq results also identified the cystine/cysteine uptake transporter TcyABC as necessary in excess L-serine, suggesting that both peptide and cysteine uptake contribute to L-serine tolerance. In addition to uptake mechanisms, glycine and diglycine toxicity is neutralized by D-alanine aminotransferase (Dat), which is required for D-alanine synthesis. The requirement for Dat and DtpT function—but not AapA—in excess glycine is explained by excess glycine inhibiting alanine uptake. Building on this finding, we found that combined treatment with glycine and the alanine analog antibiotic D-cycloserine was strongly synergistic in inhibiting *S. aureus* growth. Overall, our findings identify targetable mechanisms underlying excess amino acid tolerance in *S. aureus*, with implications for developing novel combination treatments using the accessible and biocompatible amino acids glycine and L-serine.

**IMPORTANCE:** Growing evidence supports the beneficial effects of glycine and L-serine supplementation in combating bacterial infections. Previous researchers have found that combining antibiotic treatment with high glycine concentrations has additive effects with many antibiotics, even reversing resistance to antibiotics in some bacteria. *In vivo*, activating glycine and L-serine metabolism heightens the sensitivity of bacterial pathogens to the host complement system, and studies of glycine or L-serine treatments show low toxicity and reduced inflammation in mouse and human subjects. This study reveals that glycine may be an effective antibiotic adjuvant with D-cycloserine, and treatment with glycine or L-serine could potentiate other drugs that target alanine or cysteine metabolism, respectively.

## INTRODUCTION

The amino acids glycine and L-serine have emerged as promising agents in the fight against bacterial infections. These amino acids have been used *in vivo* to treat bacterial infections as antibiotic adjuvants or alone [1–3]. Glycine and L-serine treatments have the potential to reduce morbidity and inflammation in infection [1, 2, 4]. Activation of glycine and L-serine metabolism in bacterial pathogens also renders many bacteria, including *S. aureus,* sensitive to host complement-mediated killing [3, 5]. Despite these effects, *S. aureus* requires large amounts of these amino acids during infection for protein and cell wall synthesis, making the development of resistance to glycine or L-serine treatments unlikely [6].

*In vivo* glycine and L-serine treatments to date have been effective by altering bacterial metabolism, with administered concentrations remaining well below toxic levels. Stopping bacterial proliferation requires concentrations exceeding those achievable *in vivo*, despite the low risk these amino acids pose to the host [1, 3, 7, 8]. Thus, increasing the antibacterial potency of glycine and L-serine against bacteria is necessary if they are to be used therapeutically [9, 10], and understanding the mechanisms of resistance to glycine and L-serine toxicity in bacterial pathogens may inform new treatment strategies using these amino acids. The aim of this study is to characterize genes mediating resistance to glycine and L-serine toxicity in *S. aureus,* with the goal of laying the foundation for new treatments that use glycine and L-serine to directly inhibit bacterial pathogens while also decreasing host morbidity.

Previous studies of glycine toxicity determined that excess glycine inhibits bacterial growth primarily by inhibition of cell wall synthesis through antagonism of alanine-utilizing enzymes and misincorporation in place of alanine in peptidoglycan cross-links [11, 12]. Additive or synergistic effects of glycine with cell wall-targeting antibiotics have been observed [9, 13]. Treatment with glycine has even been shown to reverse antibiotic resistance [14]. How bacteria avoid toxic accumulation of glycine is not well understood, though in *Streptomyces,* the glycine cleavage system (GCV) appears to play a significant role in detoxification [15]. Degradation of glycine by the GCV or conversion to L-serine and subsequent digestion by serine dehydrogenase represent the two major routes of glycine and L-serine catabolism in *S. aureus* [16], though the actual role of catabolism in detoxifying excess glycine or L-serine in *S. aureus* has not been studied.

L-serine toxicity has been best studied in *Bacillus subtilis,* where uptake of other amino acids (particularly threonine, arginine, aspartate or alanine) was protective [17]. Inactivation of L-serine transporters also increased resistance to L-serine, as did overexpression of threonine biosynthesis genes or L-serine catabolism genes. The current consensus is that L-serine toxicity is primarily caused by inhibition of threonine metabolism in *B. subtilis* [18].

Despite this information regarding the mechanisms of resistance to glycine and L-serine toxicity in other bacterial species, it is unclear how *S. aureus* deals with glycine and L-serine stress. To understand how *S. aureus* resists L-serine and glycine toxicity, we employed a transposon-insertion library in strain TM283, a modified strain of USA300 lineage. Results from our transposon insertion sequencing (Tn-seq) screen indicated that upon exposure to high levels of glycine or L-serine, *S. aureus* depends heavily on genes needed for uptake and synthesis of other amino acids to balance amino acid ratios disrupted by excess glycine or L-serine. Importantly, we found that excess glycine antagonizes this effort by inhibiting alanine uptake, resulting in a skewed alanine to glycine ratio as well as alanine starvation in mutants unable to synthesize alanine.

## MATERIALS AND METHODS

### Bacterial strains and growth media

Strains and plasmids used in this study are listed in Tables S1 and S2, respectively. The wild type *S. aureus* strain used for Tn-seq experiments was TM283, derived from the USA300-TCH1516 strain [19]. For making clean deletions, a nearly isogenic strain JE2 was used (99.96% average nucleotide identity with TM283). Overnight *S. aureus* cultures and sub-cultures were routinely grown in tryptic soy broth (TSB). Tryptone glucose medium (TGM) was used in all experimental cultures unless otherwise stated. This medium was adapted from that previously reported by Hussain et al. [20] and Sebulsky et al. [21]. Chemically defined medium (CDM) was also used, which differs from TGM in that individual amino acids are added in place of tryptone, and 0.2% glucose (rather than 0.1%) was added to improve growth. Both TGM and CDM formulations are detailed in the Supplemental Materials. For the process of allelic exchange, both *E. coli* and *S. aureus* were grown in brain heart infusion broth, with the addition of 10 µg/mL chloramphenicol when appropriate.

### Growth curve experiments

For growth curve experiments, *S. aureus* strain JE2 and derived mutants were prepared by growth overnight in TSB. A 1:50 dilution of the overnight culture was then sub-cultured in TSB to mid-exponential growth phase, and cells were pelleted and re-suspended in phosphate-buffered saline (PBS; 8 g/L NaCl,0.2 g/L KCl, 1.44 Na_2_HPO_4_, 0.24 KH_2_PO_4_). The suspended cells were then diluted to an optical density (OD_600_) of 0.25, and 30 µL/mL added to the experimental medium. The concentration of the inoculum was doubled when grown in defined medium to support robust growth. For growth curve experiments, 3 wells of a 96-well plate were inoculated with 150 µL of the experimental medium containing bacteria for three technical replicates, and OD_600_ was measured hourly by a BMG SpectroStar Nano plate reader with shaking at 37°C.

### Transposon insertion sequencing (Tn-seq) experiments

A transposon insertion library was created in the TM283 strain. The creation of the TM283 mutant library was accomplished using methods established by Santiago et al [22]. Phages, strains and plasmids required for this process were generously provided by Tim Meredith (Penn State University). For the experiments utilizing the transposon insertion library, a 1 mL frozen aliquot of the original library was thawed and added to 15 mL TSB. This was incubated for three hours at 37°C with shaking. 1 mL of this sub-culture was then pelleted and re-suspended in PBS, and ∼5×10^7^ CFUs (as determined by OD_600_) were transferred to experimental conditions (5 mL of TGM, TGM+125 mM glycine, TGM+500 mM diglycine or TGM+125 mM L-serine) and grown overnight with shaking at 37°C. This was done with three separate cultures of each condition. 1.5 mL aliquots were collected and DNA extracted from each culture. DNA was then processed with adapters and barcodes ligated before Illumina sequencing by SeqCenter. In processing the raw fastq file from Illumina sequencing, barcode binning, trimming and adapter removal were accomplished using the Galaxy web service [23], applying the “Barcode Splitter”, “Trim”, “Cutadapt” tools, respectively on raw fastq files. Reads were then filtered by the “Filter by Quality” tool [24, 25]. Reads were mapped back to the genome of *S. aureus* strain USA300_FPR3757 (Genbank accession: CP000255.1) using Bowtie2 [26]. 133,819 independent insertions (46 insertions per kilobase) were identified by the appearance of at least one read in TGM after stringent filtering of reads to ensure accurate mapping. Fold changes and significance of results were calculated using Bioconductor in R, where a raw P-value was produced using a moderated t-test and compared to a t-distribution. A Benjamini-Hochberg false discovery rate was then applied to get an adjusted P-value. KEGG analysis of the Tn-seq results was performed using the DAVID online web service [27]. Searching genes for encoded protein function was performed using PANTHER [28].

### *S. aureus* deletion mutant construction

Gene deletions were performed in the JE2 background (unless otherwise specified) using the pIMAY-Z plasmid and protocol from Monk et al. [29]. Briefly, PCR products (see Supplemental Materials) from flanking regions of the gene targeted for deletion were stitched together using primer overhangs and restriction digest and ligation were used for insertion of the PCR product into pIMAY-Z. The ligated plasmid + PCR insert was transformed into *E. coli* strain IM08B, and the plasmid confirmed by PCR to contain insert was isolated and transformed into the specified *S. aureus* strain. The process of allelic exchange was then performed as detailed by Monk et al. [29]. The *alr* and *aapA* transposon mutants were provided by the Nebraska Transposon Mutant Library [30].

### Glycine and L-serine MIC tests and D-cycloserine checkerboard assay

The MICs of glycine, L-serine and D-cycloserine (DCS) were determined at 37°C with shaking in TGM. Concentrations that prevented growth after 15 hours incubation were considered inhibitory. Glycine and L-serine (Sigma) were added to TGM to specified concentrations and the supplemented medium was then filter sterilized. The checkerboard assay with glycine and D-cycloserine was performed in TGM. After dilutions of the DCS and glycine were prepared, wild type JE2 or *aapA*::tn strains were inoculated at 10^6^ CFUs/mL (as measured by OD_600_) into individual wells. Inhibition was tested at 37°C with shaking, and reported results are from 15 hours of incubation.

### Mass spectrometry analysis

*S. aureus* strain JE2 was grown in TSB overnight; cells were washed and re-suspended in PBS, then inoculated into defined medium or defined medium supplemented with 100 mM glycine to a starting OD_600_ of 0.02. These cultures were grown for 8 hours, at which point cells were pelleted, supernatant filter sterilized and immediately frozen. Extractions and targeted metabolomics were performed at the BYU Metabolomics Center at Brigham Young University. Samples (500 µL) were thawed on ice and quenched with 800 µL of ice-cold methanol. 2 µL of internal standard was added (MSK-QRESS1-1, Cambridge Isotope Laboratories, Inc., Tewksbury, MA) followed by a 30-minute incubation at -80°C. After re-thawing and vortexing, the suspension was centrifuged at 20,000 g for 20 minutes at 4°C. The resulting supernatant was transferred to a glass vial, evaporated to dryness under a N2 stream at 30°C, and reconstituted in 120 µL of ice-cold acetonitrile:water (60:40, v/v). The reconstituted mixture was centrifuged (20,000g, 10 min, 4°C) and the supernatant filtered using a 0.2µm centrifuge filter for 2 minutes (300 rpm, 4°C). 2 µL from the bottom aqueous layer of the resulting biphasic sample was diluted with 198 µL of 60:40 acetonitrile:water in a glass insert for LC-MS analysis. Samples were run on a Thermo Orbitrap Exploris 480 mass spectrometer coupled to a Vanquish Horizon liquid chromatography system with an iHILIC-(P) Classic HILICON AB column (2.1×150 mm, 5 μm) in negative mode with 20mM bicarbonate in water (pH=9.6) and acetonitrile using an analytical method described by Sing et al., (2024). Raw acquisition data was processed using a targeted metabolomics workflow in the Skyline metabolomics application and amino acids were identified and quantified using authentic standards and calibration curves (R^2^ > 0.99), respectively.

### Statistical analysis

For statistical analysis of Tn-seq results, a raw P-value was produced using a moderated t-test and compared to a t-distribution using Bioconductor in R. A Benjamini-Hochberg false discovery rate was then applied to get an adjusted P-value. Other statistical analyses were performed using GraphPad Prism 10 software (GraphPad Software, San Diego, CA, USA). Statistically significant differences between two groups were established by an unpaired t-test. Growth curve experiments are reported as the mean of three separate experiments, with error bars showing the standard deviation between experiments.

## RESULTS

### Tn-seq identifies genes mediating resistance and susceptibility to amino acid toxicity

Excess glycine causes toxicity in *S. aureus* mainly by its misincorporation in place of alanine by cell wall-biogenesis enzymes [11, 12]. Diglycine (a dipeptide of glycine, gly-gly) may be toxic by the same mechanisms, but require different transporters and catabolism into monomers before toxicity occurs. L-serine toxicity has not been characterized in *Staphylococcus,* but may inhibit threonine metabolism, as seen in *Bacillus* [18]. To determine the genes and associated processes that are essential for growth in the presence of high but sub-inhibitory concentrations of L-serine, glycine, or diglycine, a transposon insertion sequencing (Tn-seq) experiment was performed using a library created in strain TM283. This experiment was performed in 125 mM L-serine (MIC = 500 mM), 125 mM glycine (MIC = 500 mM), or 500 mM diglycine (MIC = >1 M). The library cultures were grown to saturation in tryptone glucose medium (TGM), which provides both peptides and free amino acids. Genes were ranked based on the fold-change (FC) in gene insertion counts between treatment and control populations. A negative FC value indicates depletion of counts in the treatment condition and therefore conditional essentiality, while a positive FC value indicates the functional gene is deleterious in the treatment condition. Volcano plots of the results of these experiments are shown in Figure 1A-C. Genes with a Log_2_FC < -3 and adjusted P-value < 0.05 are considered conditionally essential in the treatment condition. As this study focuses on individual testing of conditionally essential genes and not those that are detrimental, we expanded our definition of “deleterious” genes to those with a Log_2_FC > 2 (rather than 3) and adjusted P-value < 0.05 to provide a broader overview of potentially deleterious pathways.

**Figure 1:**
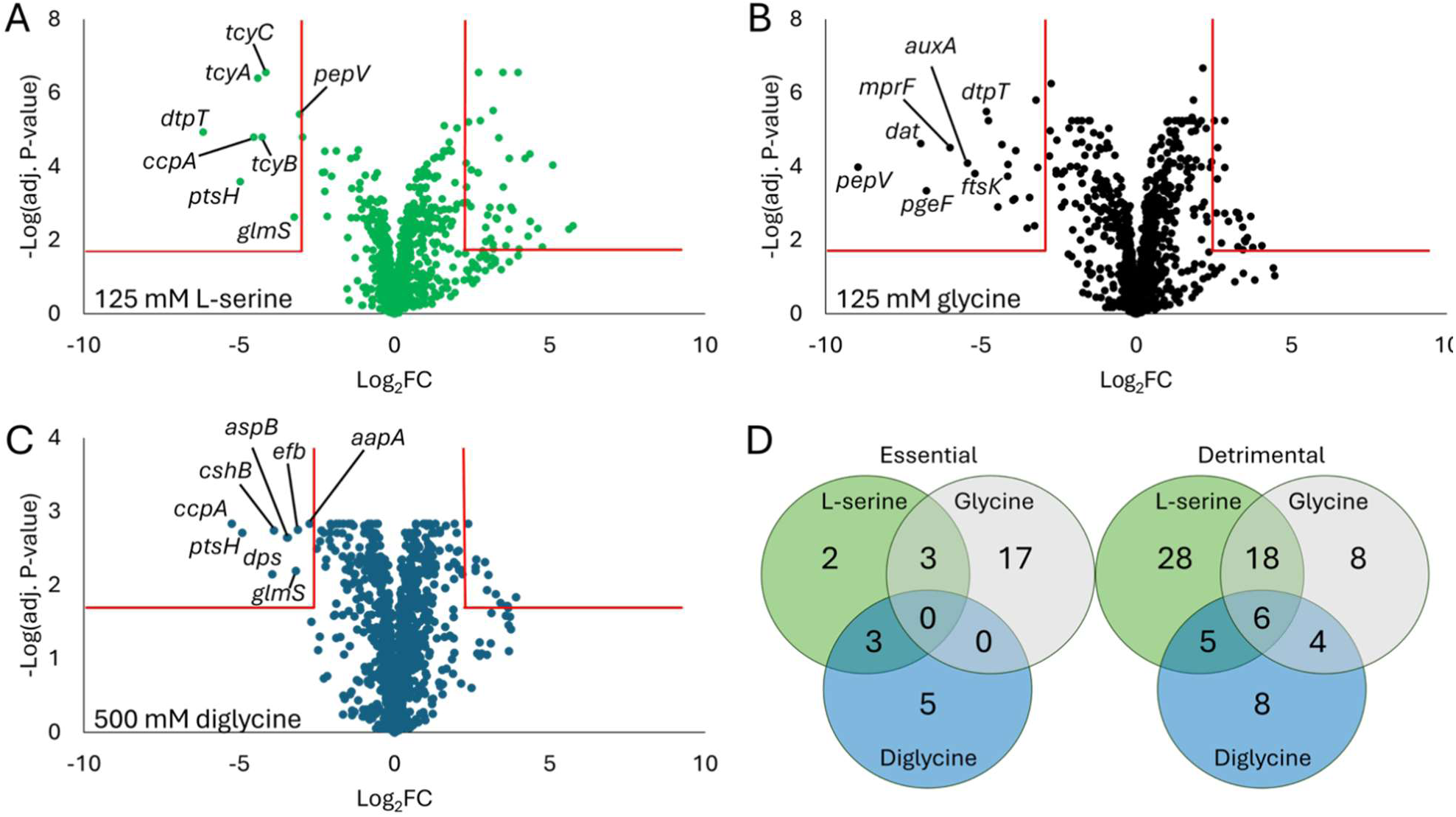
Volcano plots of transposon insertion sequencing experiments show conditionally essential or deleterious genes in excess L-serine, diglycine and glycine. **(A)** Volcano plot of *S. aureus* strain TM283 transposon insertion library grown in 125 mM L-serine, showing Log_2_(Fold Change (FC) in reads mapping back to the gene) on the x-axis and -Log(adjusted P-value) on the y-axis; red lines mark the constraints defining conditionally essential genes (Log_2_FC < -3; adjusted P-value < 0.05; genes found in the upper left) or conditionally detrimental (Log_2_FC > 2; adjusted P-value < 0.05; genes found in the upper right). **(B)** Volcano plot of *S. aureus* transposon insertion library grown in 500 mM diglycine. **(C)** Volcano plot of *S. aureus* transposon insertion library grown in 125 mM glycine. Tn-seq experiments were each performed in triplicate. For statistical analysis of Tn-seq results, a raw P-value was produced using a moderated t-test and compared to a t-distribution. A Benjamini-Hochberg false discovery rate was then applied to get an adjusted P-value.

For growth in excess L-serine, 8 genes were conditionally essential, and 58 genes were detrimental. For glycine, 20 genes were conditionally essential, and 36 were detrimental. For diglycine, 8 genes were conditionally essential, and 23 were detrimental. Overlap in conditionally essential genes between conditions was limited, and no genes were essential in all three treatments. In contrast, 6 genes were found to be detrimental in every treatment condition (Figure 1D).

### The TCA cycle and oxidative respiration potentiate glycine and L-serine toxicity

KEGG analysis of genes that were deleterious in all three amino acid treatments showed that they predominantly play roles in the tricarboxylic acid (TCA) cycle (L-serine treatment q-value = 3.32e-2; glycine treatment q-value = 4.09e-4; diglycine treatment q-value = 2.62e-5) and oxidative phosphorylation (L-serine treatment q-value = 4.78e-11; glycine treatment q-value = 1.82e-3; diglycine treatment q-value = 5.84e-3). Succinate dehydrogenase, which converts succinate to fumarate in the TCA cycle, was deleterious in every treatment (average Log_2_FC of *sdhABC* genes in L-serine = 3.11; in glycine = 3.40; in diglycine = 3.67). Fumarate hydratase, which converts fumarate to malate, also had negative effects on growth across all treatments (Log_2_FC of *fumC* in in L-serine = 3.23; glycine = 3.68; in diglycine = 3.67). Specific genes involved in oxidative respiration were not shared as consistently, though genes from that pathway were highly represented in all treatments. Considering these results together, it appears that active cellular respiration (TCA cycle or electron transport) causes increased susceptibility to excess glycine or L-serine.

In predicting other potentially deleterious processes in excess glycine and L-serine, it seemed likely that glycine or L-serine-specific transporters would facilitate toxic accumulation of glycine or L-serine, as reported in a study of amino acid toxicity in *Bacillus* [18]. In our Tn-seq experiment in excess glycine, two amino acid transporters were identified with detrimental functions: the glycine betaine transporter gene *opuD* (Log_2_FC = 2.14) and an uncharacterized amino acid transporter SAUSA300_1231 (Log_2_FC = 2.09). No amino acid or peptide transporter genes were identified as deleterious in diglycine or L-serine treatments. The absence of an exceptionally deleterious transporter in any treatment condition may indicate functional redundancy of multiple transporters, or that high concentrations of these amino acids cause transport to occur non-specifically.

### Genes essential in excess L-serine

The eight genes conditionally essential for *S. aureus* growth in 125 mM L-serine are shown in Table 1. Mutation of the *dtpT* gene encoding a dipeptide transporter had the strongest effect on fitness upon L-serine treatment (Log_2_FC = -6.16). The DtpT transporter has low specificity in the amino acid composition of peptides it transports, though it is fairly specific to dipeptides [31–33]. DtpT has some redundant function to other transporters such as Opp3 (Log_2_FC = 0.9)[34], making the importance of DtpT noteworthy given the highly diverse peptides and free amino acids available in the tryptone supplied by TGM [35]. Monoculture testing of an in-frame deletion mutant of *dtpT* in 125 mM L-serine confirmed its enhanced sensitivity (Figure 2A). The importance of peptide uptake by DtpT may be to supply various amino acids that offset the consequences of intracellular L-serine accumulation, as observed in *Bacillus* [17, 18]. However, the peptides taken up by DtpT must be hydrolyzed to free amino acids before metabolic use, and this may explain the importance in excess L-serine of a putative dipeptidase encoded by the *pepV* gene (Log_2_FC = -3.06). The *pepV* gene is part of a bicistronic operon, located upstream of the *dat* gene (Log_2_FC = 0.06), which encodes a D-alanine aminotransferase that fulfills *de novo* D-alanine synthesis. Thus, transposon insertions in *pepV* may impact function of the downstream *dat* gene. We therefore made in-frame deletions of either *dat* or *pepV* genes to discern the importance of each in excess L-serine. Monoculture testing confirmed a growth defect of the *pepV* deletion mutant and not a *dat* mutant in 125 mM L-serine, validating the notion that PepV-mediated peptide hydrolysis is critical for growth in excess L-serine, independent of effects on *dat* expression (Figure 2B).

**Table 1.**
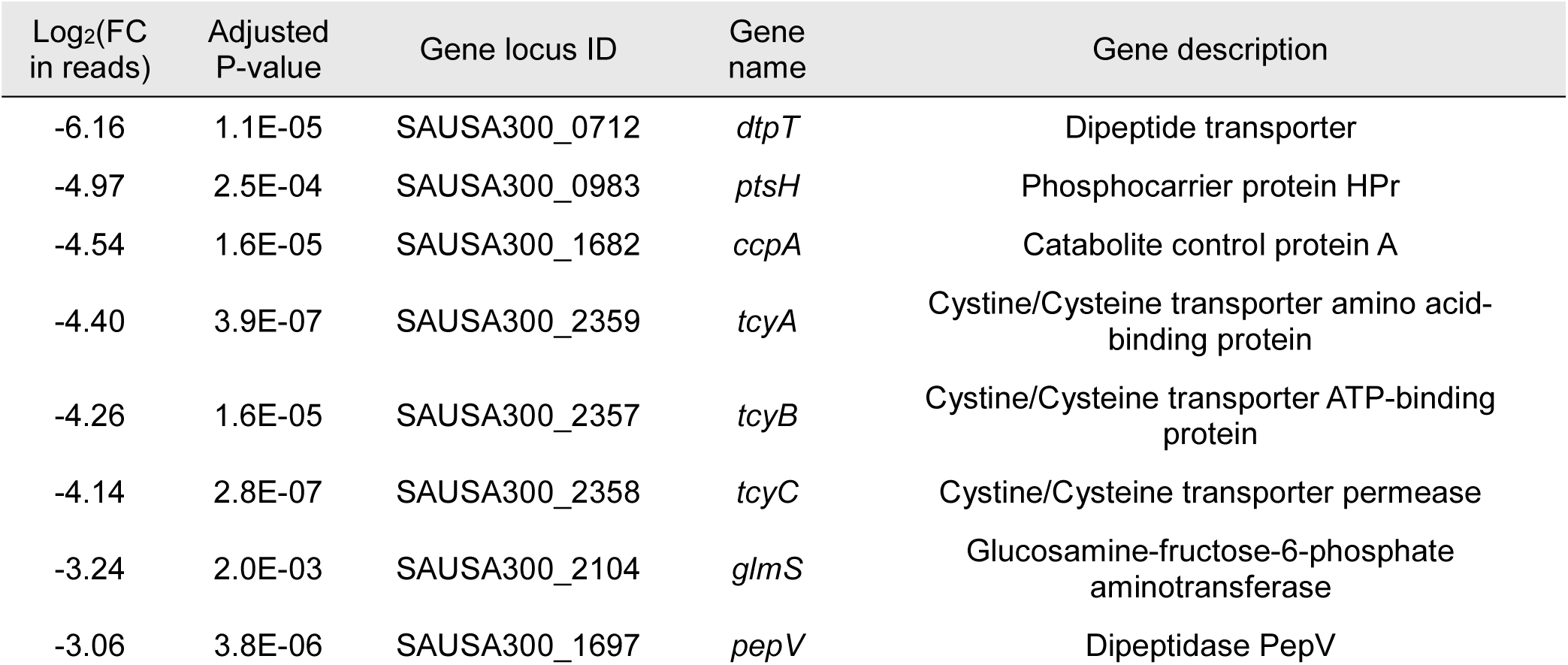
Genes essential for fitness in 125 mM L-serine.

**Figure 2:**
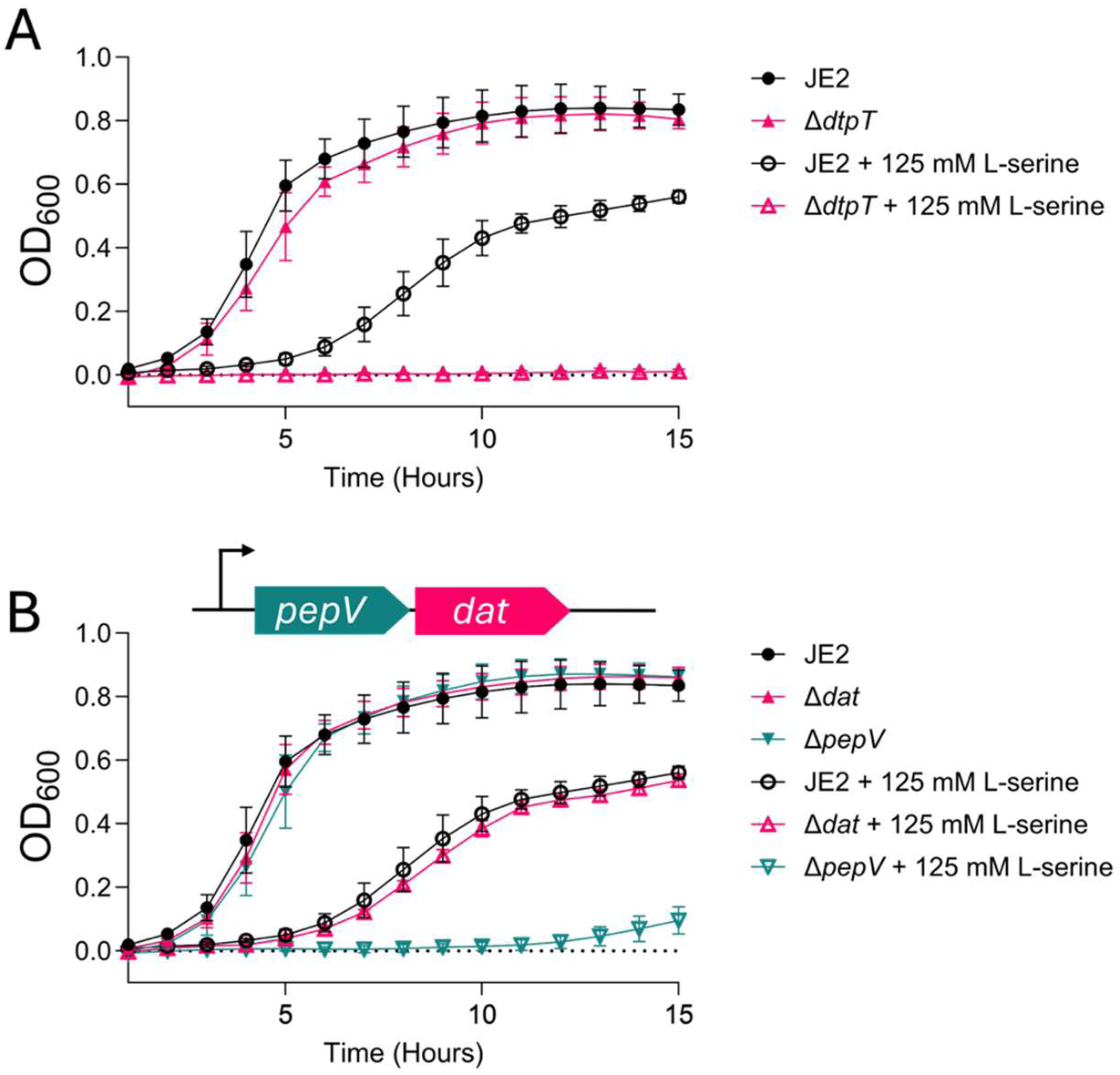
Peptide uptake by DtpT and peptide catabolism by PepV are necessary for growth in excess L-serine. **(A)** Growth of a deletion mutant of the *dtpT* gene in TGM or TGM containing 125 mM L-serine. **(B)** Growth of deletion mutants of *dat* or *pepV* in TGM or TGM containing 125 mM L-serine. Growth curve experiments are reported as an average of three separate experiments, performed in triplicate. Error bars represent the standard deviation between experimental replicates.

The *ptsH* and *ccpA* genes were also important for growth in L-serine (Log_2_FC = -4.97 and Log_2_FC = -4.54, respectively). Both function in carbon catabolite repression and glucose utilization, with HPr (encoded by *ptsH*) sensing glucose availability and acting as an allosteric activator of CcpA, resulting in positive regulation of glucose-utilizing genes and negative regulation of alternative carbon source catabolism, including amino acids [36, 37]. The essentiality of CcpA and HPr suggests that L-serine toxicity is exacerbated by glucose starvation. This may be caused by glucose starvation stimulating digestion of amino acids that offset L-serine toxicity, or an increase in L-serine uptake upon glucose starvation [16].

The *tcyABC* genes, encoding subunits of a single cystine/cysteine uptake transporter, were also among the conditionally essential genes in excess L-serine. *S. aureus* produces two L-cysteine transporters, encoded by *tcyABC* (average Log_2_FC = -4.27), and *tcyP,* which did not have a notable effect on L-serine tolerance (Log_2_FC = 0.13). Consistent with these results, previous work has shown that TcyABC is the functionally active L-cysteine transporter in culture [38]. The expression of *tcyABC* is controlled by the CymR repressor [39], and unsurprisingly, *cymR* was identified as a deleterious gene in the Tn-seq experiment with excess L-serine (Log_2_FC = 4.27). CymR also represses L-cysteine synthesis in addition to uptake, controlling the transcription of genes in the methionine-dependent synthesis pathway as well as the L-serine-dependent pathway [39]. Curiously, *cysE*, encoding the serine acetyltransferase necessary for L-serine-dependent cysteine synthesis, had a deleterious effect on fitness upon L-serine treatment (Log_2_FC = 3.06). How L-cysteine uptake and not biosynthesis protects against L-serine toxicity is unclear. It is possible that L-serine or O-acetylserine (synthesized by CysE using L-serine) accumulation is toxic or disrupts L-cysteine synthesis, rendering TcyABC-dependent uptake of L-cysteine essential. This potential competition between L-cysteine and L-serine in L-cysteine metabolism may relate to the structural similarity between these two amino acids.

When predicting essential genes in excess L-serine, it seemed likely that the serine-catabolizing activity of serine dehydrogenase would be protective against intracellular accumulation. The *sdaAA* and *sdaAB* genes encode the alpha and beta subunits of serine dehydrogenase, respectively. Our Tn-seq data indicate a modest fitness defect upon loss of these genes in excess L-serine (Log_2_FC of *sdaAA* = -1.11; Log_2_FC of *sdaAB* = -1.22). Consistent with this, a double-deletion mutant was created and tested in monoculture for sensitivity to L-serine, and found to grow slightly slower than the wild type (Figure S1A). We concluded that serine dehydrogenase activity does not strongly influence L-serine toxicity.

### Genes essential in excess glycine

The 20 conditionally essential genes in 125 mM glycine are listed in Table 2. Transposon insertions in the genes *pepV* and *dat* (located in the same operon) resulted in the strongest fitness defects (Log_2_FC of *pepV* = -8.96; Log_2_FC of *dat* = -6.94). In monoculture, a *dat* deletion mutant was unable to grow in glycine concentrations above 80 mM, while a *pepV* deletion mutant showed no difference in glycine sensitivity relative to wild type JE2 (Figure 3A). Given that *pepV* is upstream of *dat* in this operon, the apparent essentiality of *pepV* in the Tn-seq dataset is likely due to polar effects on Dat expression caused by transposon insertions in *pepV*. Thus, the strong Tn-seq result associated with *pepV* most likely reflects the importance of Dat activity. We propose that Dat is required to supply D-alanine to counterbalance high glycine accumulation, competitively preventing misuse of glycine by alanine-dependent cell wall-biogenesis enzymes inside the cell.

**Table 2.**
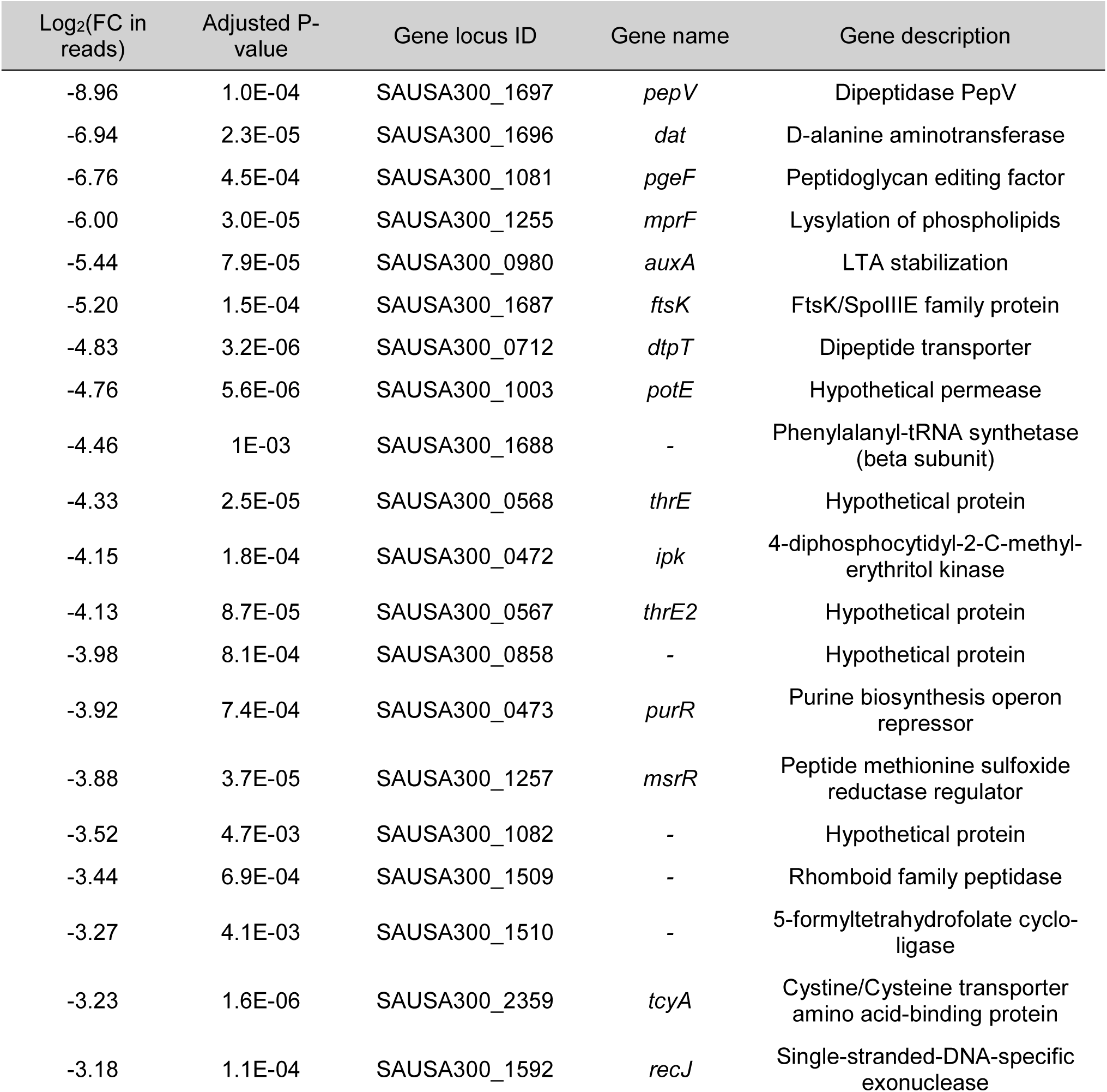
Genes essential for fitness in 125 mM glycine.

**Figure 3:**
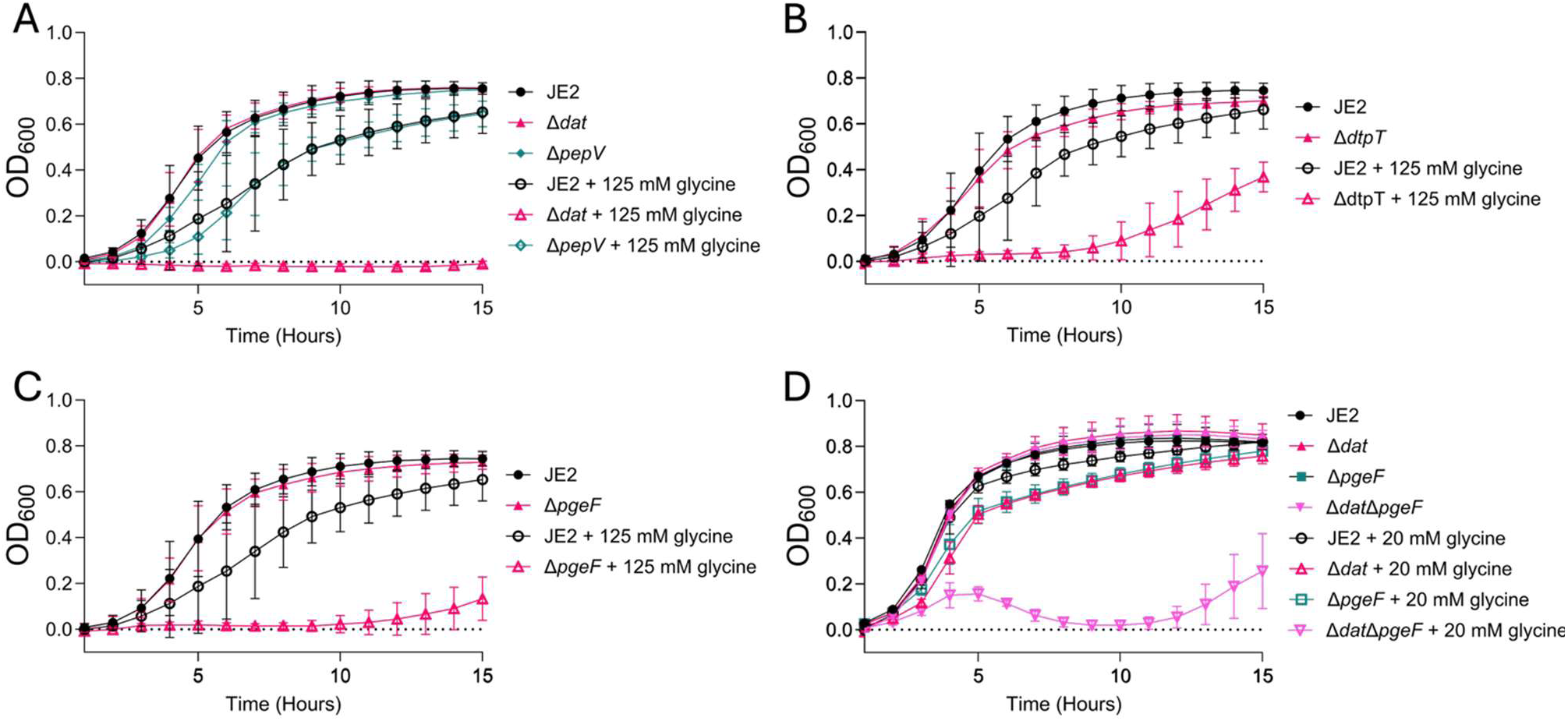
Alanine synthesis by Dat, peptide uptake by DtpT, and peptidoglycan cross-link editing by PgeF are critical processes for growth in excess glycine. **(A)** Growth of deletion mutants of the alanine synthesis gene *dat* or the putative dipeptidase gene *pepV* in TGM or TGM containing 125 mM glycine. **(B)** Growth of a deletion mutant of the dipeptide transporter gene *dtpT* in TGM or TGM containing 125 mM glycine. **(C)** Growth of a deletion mutant of the peptidoglycan editing factor gene *pgeF* in TGM or TGM containing 125 mM glycine. **(D)** Growth of a double deletion mutant of *dat* and *pgeF* grown in TGM or TGM containing 20 mM glycine. Growth curves are reported as an average of three separate experiments, performed in triplicate. Error bars represent the standard deviation between experimental replicates.

The dipeptide transporter gene *dtpT* was also important in excess glycine (Log_2_FC = -4.83), and growth of a *dtpT* deletion mutant in monoculture confirmed its heightened sensitivity (Figure 3B). As reported above, the DtpT dipeptide transporter is also required for L-serine tolerance. Like in excess L-serine, DtpT-mediated peptide uptake may be beneficial in excess glycine because it supplies amino acids to favorably compete with glycine. Considering the demonstrated importance of DtpT-mediated dipeptide uptake in subverting glycine toxicity, it is notable that the PepV dipeptidase—required for L-serine tolerance—does not appear important for glycine tolerance (Figure 3A). This difference may reflect substrate specificity of PepV, such that it preferentially releases amino acids that mitigate L-serine toxicity but not glycine toxicity.

Cell envelope maintenance and integrity genes were also conditionally essential upon glycine treatment. The gene SAUSA300_1081 (Log_2_FC = -6.76) encodes an ortholog of the *pgeF* gene in *E. coli,* a peptidoglycan-editing amidase that removes erroneously incorporated amino acids (such as glycine and L-serine) from peptidoglycan cross-links [40]. The function of *pgeF* orthologs in gram-positive bacteria have been shown to be broadly conserved [41], indicating that PgeF of *S. aureus* performs a similar role. The necessity of cross-link editing in excess glycine is consistent with the established mechanism of glycine toxicity, namely the misincorporation of glycine in place of alanine in peptidoglycan cross-links [12]. In monoculture, a *pgeF* deletion mutant exhibits an MIC reduced to 250 mM glycine (compared to 500 mM of the wild type) in TGM, as well as growth defects at lower concentrations (Figure 3C). We predicted that deficient D-alanine supply via a *dat* mutation would aggravate the glyine sensitivity observed in the *pgeF* mutant due to rampant glycine misincorporation in peptidoglycan. Consistent with this, deletion of *pgeF* in a *dat* deletion background resulted in extreme sensitivity to glycine, with severe growth defects of the double mutant in 20 mM glycine (Figure 3D).

That this double mutant is much more sensitive to glycine than either single mutant is consistent with the hypothesis that D-alanine synthesis and cross-link editing cooperate as independent functions to protect against glycine toxicity. This also adds to the evidence reported previously that glycine misincorporation in place of alanine in peptidoglycan is the primary mechanism of glycine toxicity in *S. aureus.* Other cell envelope-stabilizing genes identified as essential in excess glycine include *auxA* (Log_2_FC = -5.4) and *mprF* (Log_2_FC = -6.00). These two genes affect underlying cell envelope properties by stabilizing teichoic acids and neutralizing the cell membrane, respectively [42, 43]. Thus, direct removal of glycine in cross-links appears to be carried out by PgeF, and stabilization to tolerate glycine misincorporation is likely fulfilled by AuxA and MprF.

Two additional genes required for glycine tolerance are annotated as putative efflux transporters: SAUSA300_0567 (Log_2_FC = -4.13) and SAUSA300_0568 (Log_2_FC = -4.33). These genes are co-transcribed and both encode proteins that share conserved domains (PF12821 and PF06738) with ThrE, a threonine efflux transporter characterized in *Corynebacterium* [44]. This shared domain provides some evidence that these proteins may function in amino acid efflux. However, the overall length and amino acid sequence between these *S. aureus* proteins and the *Corynebacterium* homolog are limited in similarity (<=20% sequence identity), and a similar operon within the *S. aureus* genome also encodes proteins with closely related domains, though their function has no impact on glycine sensitivity (SAUSA300_0728 and SAUSA300_0729, Log_2_FC of -0.07 and 0.02, respectively). It is possible that SAUSA300_0567 and SAUSA300_0568 encode amino acid efflux transporters, and given the beneficial effects of these transporters in tolerating glycine, they may export glycine. However, the actual function of these putative transporters remains unclear.

We expected that the glycine cleavage (GCV) system would be critical based on its capacity to catabolize glycine. There is also precedence for its importance in high-glycine tolerance in *Streptomyces* [15]. However, our Tn-seq experimental results did not indicate that the GCV system is necessary upon glycine treatment. The first step in the GCV pathway is carried out by the two-subunit GcvP decarboxylase, encoded by the co-transcribed genes *gcvPA* and *gcvPB* (*gcvPA* Log_2_FC = -0.19; *gcvPB* Log_2_FC = -0.23). In monoculture testing of a *gcvPA/PB* double deletion mutant, we observed gradual lysis (based on decreasing optical density over time) upon entry into stationary phase in excess glycine (Figure S1B), but no change in MIC. The first step in the second possible route of glycine catabolism is conversion of glycine to L-serine by GlyA for eventual entry into central metabolism as pyruvate. To assess the importance of this other potential route of glycine catabolism, we created a *glyA* deletion mutant, which showed no defect in excess glycine or L-serine. This indicates that interconversion by GlyA does not assist wild type bacteria in tolerating excess glycine or L-serine (data not shown). Thus, catabolism of glycine does not appear to strongly influence susceptibility of *S. aureus* to excess glycine.

### Genes essential in excess diglycine

*S. aureus* is less sensitive to diglycine compared to glycine, with an MIC >1 M. However, treatment with >250 mM diglycine does result in a decrease in final culture turbidity (Figure S2), providing evidence of stress. Following growth of the transposon library in 500 mM diglycine, we identified eight conditionally essential genes (Table 3).

**Table 3.**
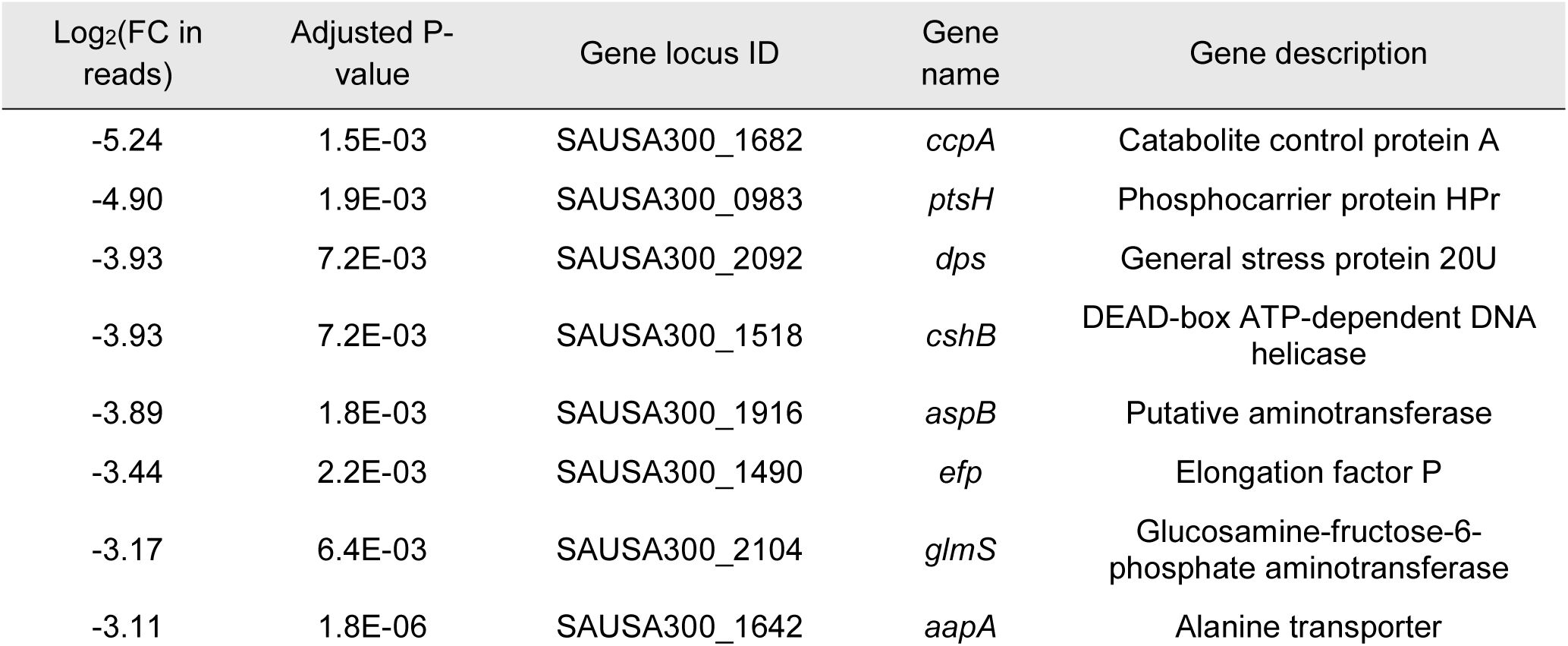
Genes essential for fitness in 500 mM diglycine.

The *ccpA* (Log_2_FC = -5.24) and *ptsH* (Log_2_FC = -4.90) genes encoding key regulators of energy and carbon metabolism have the greatest effect on fitness in diglycine. These genes were also identified as necessary for growth upon L-serine treatment (Figure 1, Table 1). We hypothesize that upregulation of peptide uptake systems occurs in *ccpA* and *ptsH* mutants as a starvation response, causing rapid accumulation and subsequent toxicity [16, 36, 37, 45]. These results provide evidence that the toxicity of diglycine and L-serine, but not glycine, is impacted by the capacity of the cell to utilize glucose.

In contrast to the results with glycine, the *dat* gene was not found to be essential in excess diglycine (log_2_FC of 0.4 in diglycine compared to -6.94 in glycine). However, we performed monoculture testing of the *dat* deletion mutant in diglycine and observed gradual lysis of the *dat* mutant cells following entry into stationary phase, with total loss of optical density by 15 hours (Figure 4). Thus, it appears that alanine synthesis does indeed play a role in resisting toxicity of both glycine and diglycine, though in diglycine the consequence of the *dat* mutation is evident only in later stages of growth. The *aapA* gene encoding an alanine uptake transporter was also among the conditionally essential genes in excess diglycine (Log_2_FC = -3.11). This is consistent with the general need for intracellular alanine to offset the effects of intracellular glycine accumulation. However, the *aapA* gene did not appear essential in free glycine (Log_2_FC = -0.51), while the dipeptide uptake transporter encoded by *dtpT*—which we posit is critical to supply alanine-containing peptides—is essential in excess glycine (Log_2_FC in glycine = -4.83) but not diglycine (Log_2_FC = 0.98). In the following section, we resolve the mechanisms behind these puzzling results by examining how free glycine treatment impacts alanine uptake and metabolism.

**Figure 4:**
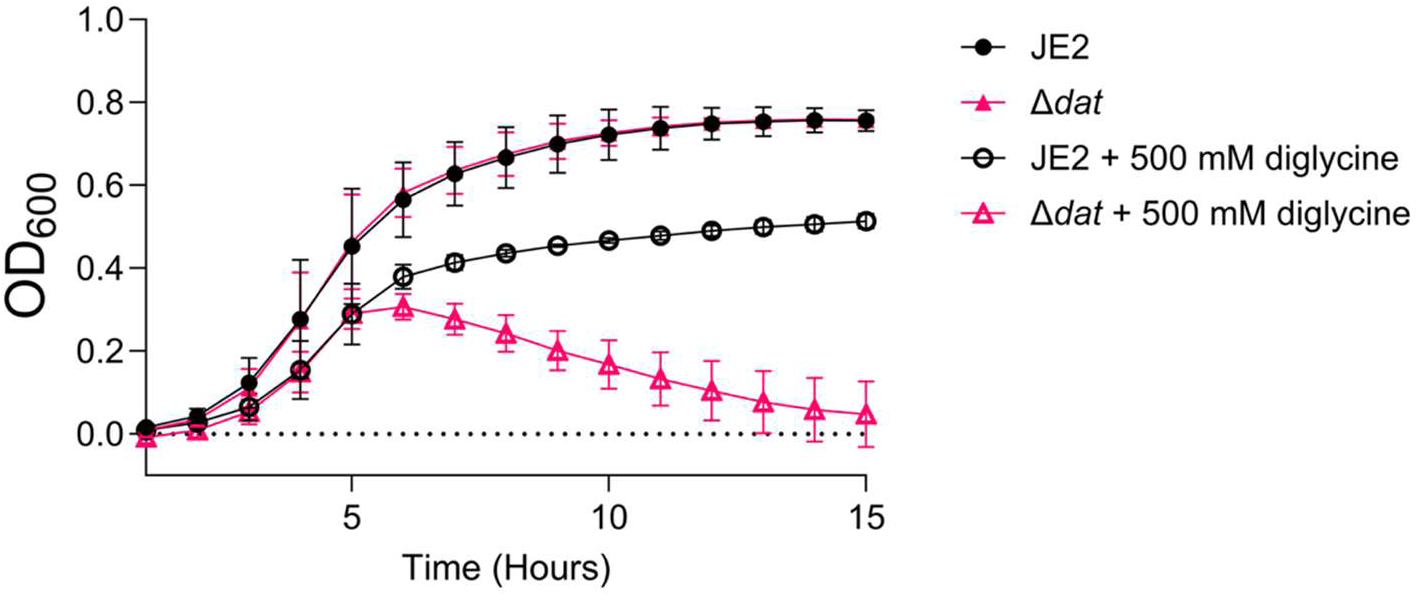
Alanine synthesis by Dat prevents lysis after entry into stationary phase when grown in 500 mM diglycine. Growth of the *dat* deletion mutant in TGM or TGM containing 500 mM diglycine. Growth curves are reported as an average of three separate experiments, performed in triplicate. Error bars represent the standard deviation between experimental replicates.

### Excess glycine inhibits alanine uptake

Under normal growth conditions in *S. aureus*, L-alanine is imported by transporters such as AapA, and can then be converted to D-alanine by the racemase Alr, which interconverts L-alanine and D-alanine [46, 47]. If exogenous L-alanine were contributing significantly to the D-alanine pool, we would expect Alr activity to be important for growth of *S. aureus* in excess glycine. This expectation is based on the mechanism of glycine toxicity and the observed contributions of D-alanine synthesis by Dat in alleviating toxicity. Yet, *alr* was not among the essential genes in our screen (Log_2_FC = -1.2), and monoculture testing confirmed that an *alr* mutant (*alr::*tn) was not more sensitive than wild type to glycine in TGM (Figure 5A). We hypothesized that this reflects an insufficient supply of L-alanine (via uptake) to support consequential conversion to D-alanine. In considering the reason for this deficient supply, the importance of peptide uptake by DtpT but unimportance of AapA-mediated uptake of free alanine in these conditions suggests that peptides—not free alanine—are serving as the main source of L-alanine under glycine stress. We predicted that this was due to excess glycine inhibiting uptake of free alanine. If glycine does indeed inhibit free alanine uptake, then in a medium containing only free amino acids, Alr activity would become necessary for endogenously supplying L-alanine by converting Dat-synthesized D-alanine to L-alanine. Consistent with this expectation, the *alr*::tn strain exhibited a growth defect relative to wild type upon excess glycine treatment in chemically defined medium (CDM), in which tryptone was replaced by free amino acids (Figure 5B).

**Figure 5:**
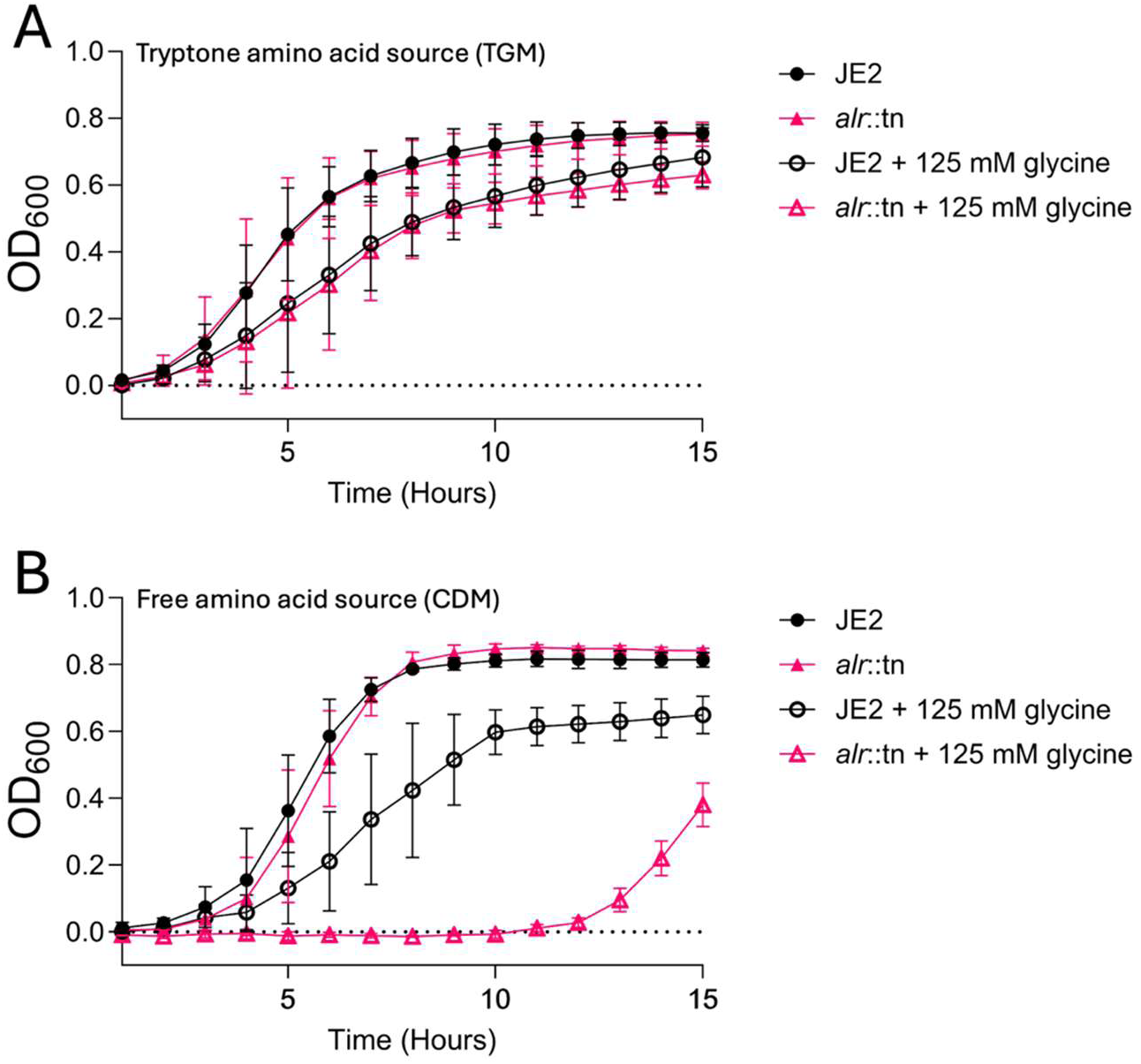
Peptides as an amino acid source in culture medium remove the need for the alanine racemase activity of Alr in excess glycine. **(A)** Growth of the *alr* transposon mutant (*alr::*tn) in TGM (tryptone amino acids source) or TGM containing 125 mM glycine. **(B)** Growth of *alr::*tn in CDM (free amino acid source) or CDM with 125 mM glycine. Growth curves are reported as an average of three separate experiments, performed in triplicate. Error bars represent the standard deviation between experimental replicates.

To provide further evidence that free alanine uptake is inhibited by excess glycine, we measured alanine depletion from the supernatant of wild-type *S. aureus* grown in CDM (this basal medium contains 1 mM L-alanine and 1 mM glycine), with and without treatment of 100 mM additional glycine. This depletion is reflective of the quantity of each amino acid consumed by the cultured cells. The growth of the untreated samples and those treated with 100 mM glycine at 8 hours was not statistically differerent (Figure 6A). As predicted, glycine treatment resulted in a 3.2-fold reduction in alanine depleted (and by extension, consumed by the cells) from the culture supernatant (Figure 6B).

**Figure 6:**
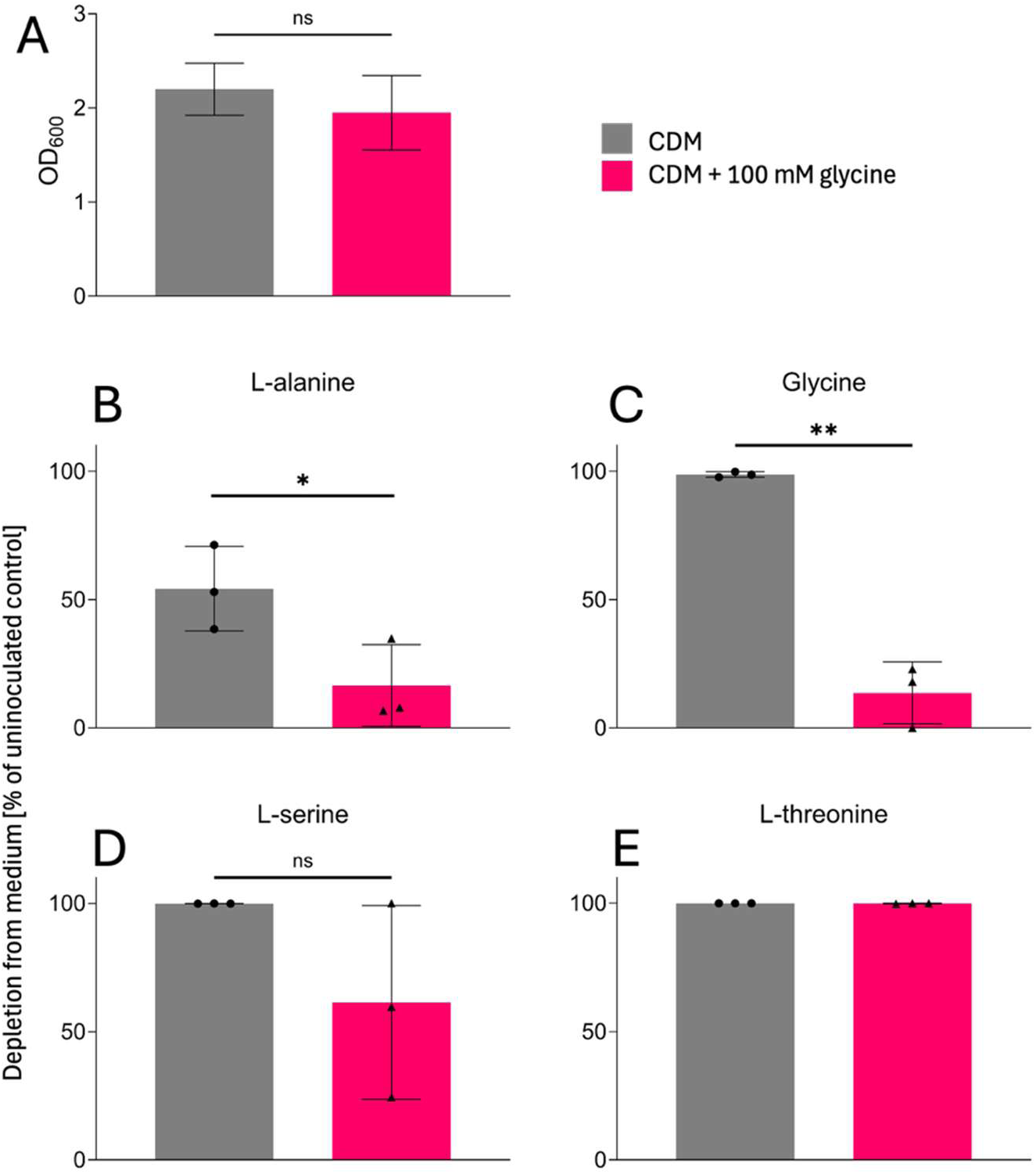
Alanine depletion from culture supernatants is reduced when excess glycine is present in the medium. **(A)** Optical density of the cultures used for LC-MS analysis after 8 hours culture in CDM at 37°C with shaking. **(B)** Measurement of alanine depleted from culture supernatants after 8 hours, reported as percent (%) depletion of the 1 mM present in CDM. **(C)** Measurement of glycine depleted from culture supernatants, reported as % depletion; in the untreated sample, this % depletion refers to the 1 mM present in basal CDM, but in the 100 mM treated sample, % depletion refers to glycine in CDM plus the 100 mM treatment. **(D)** Measurement of serine depleted from culture supernatants, reported as % depletion of the 1 mM in CDM. **(E)** Measurement of threonine depleted from culture supernatants reported as % depletion of the 1.5 mM in CDM. Reported growth and amino acid analyses were performed on three independent cultures in two separate experiments. Statistical significance was determined using an unpaired, two sample t-test (* = P-value < 0.05; ** = P-value < 0.005).

Antagonism of alanine uptake may be caused by the high amount of glycine that remains in the culture medium (Figure 6C). Glycine treatment may also impact L-serine uptake (Figure 6D), while having no effect on total depletion of L-threonine from the medium (Figure 6E).

Overall, these findings indicate that excess glycine, even at sub-inhibitory concentrations, prevents the uptake of L-alanine by *S. aureus.* This likely explains the essentiality of Dat-mediated alanine synthesis in excess glycine, and explains the importance of peptide uptake as a supply of exogenous L-alanine when cells are grown in TGM with excess glycine. Our interpretation of these results is summarized diagramatically in Figure 7.

**Figure 7:**
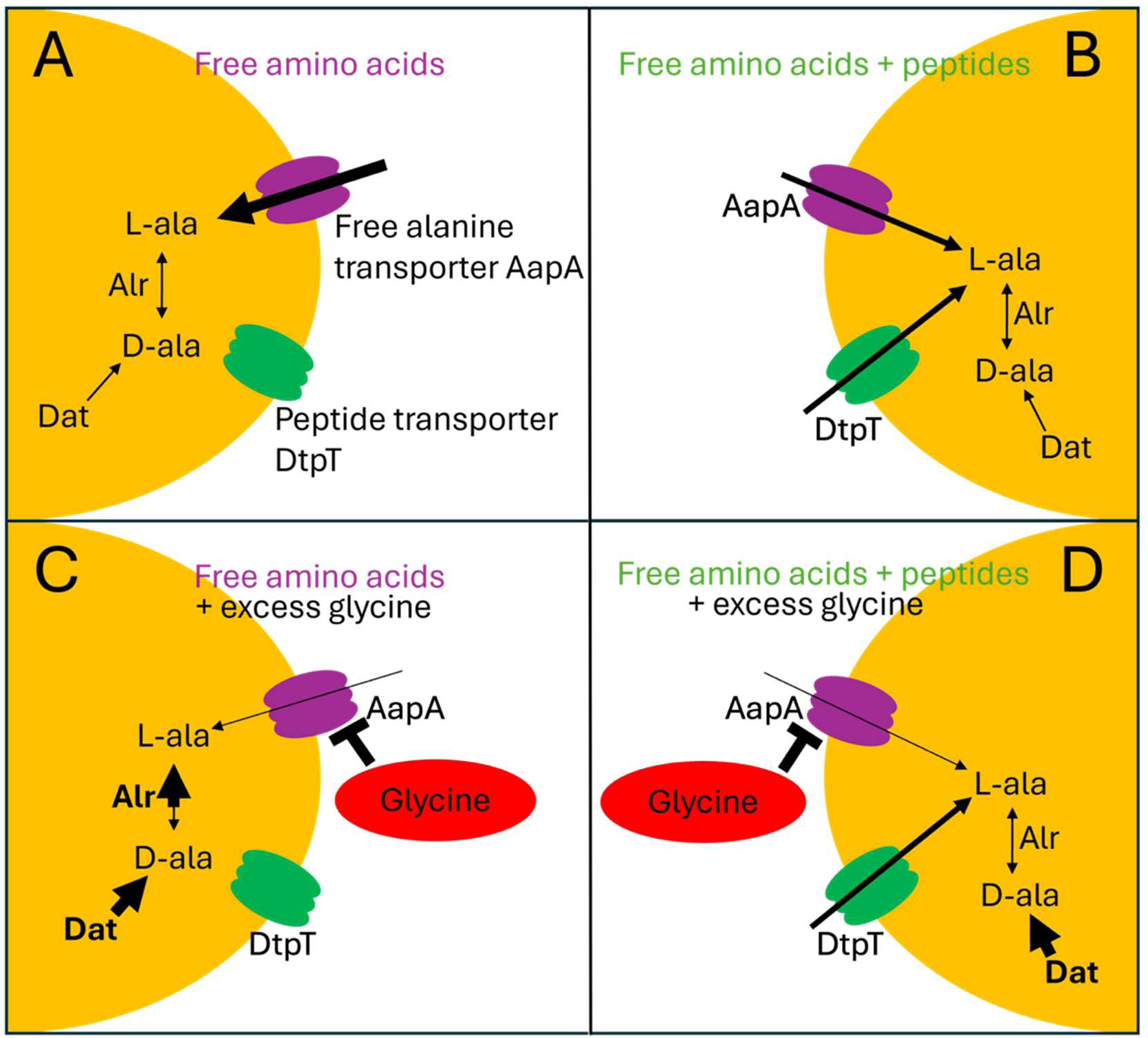
A depiction of the proposed effect of excess glycine on alanine metabolism. Processes that are heavily utilized or essential in the given condition are in bold. **(A)** Alanine acquisition in the presence of exclusively free amino acids (as in CDM) requires free amino acid transporters such as the alanine transporter AapA. **(B)** Alanine acquisition in the presence of both free amino acids and peptides (as in TGM) uses both free amino acid transporters and peptide transporters, such as the dipeptide transporter DtpT. **(C)** Growth in excess glycine with only free amino acids available (as in CDM) requires Dat activity to counterbalance glycine toxicity, and Dat and Alr to supply L-ala because of glycine-mediated inhibition of free alanine uptake. **(D)** Growth in excess glycine with free amino acids and peptides available reverses the necessity of Alr because of the exogenous supply of alanine *via* peptides, but Dat activity is still necessary to supply D-alanine to counterbalance glycine toxicity.

**Figure 8:**
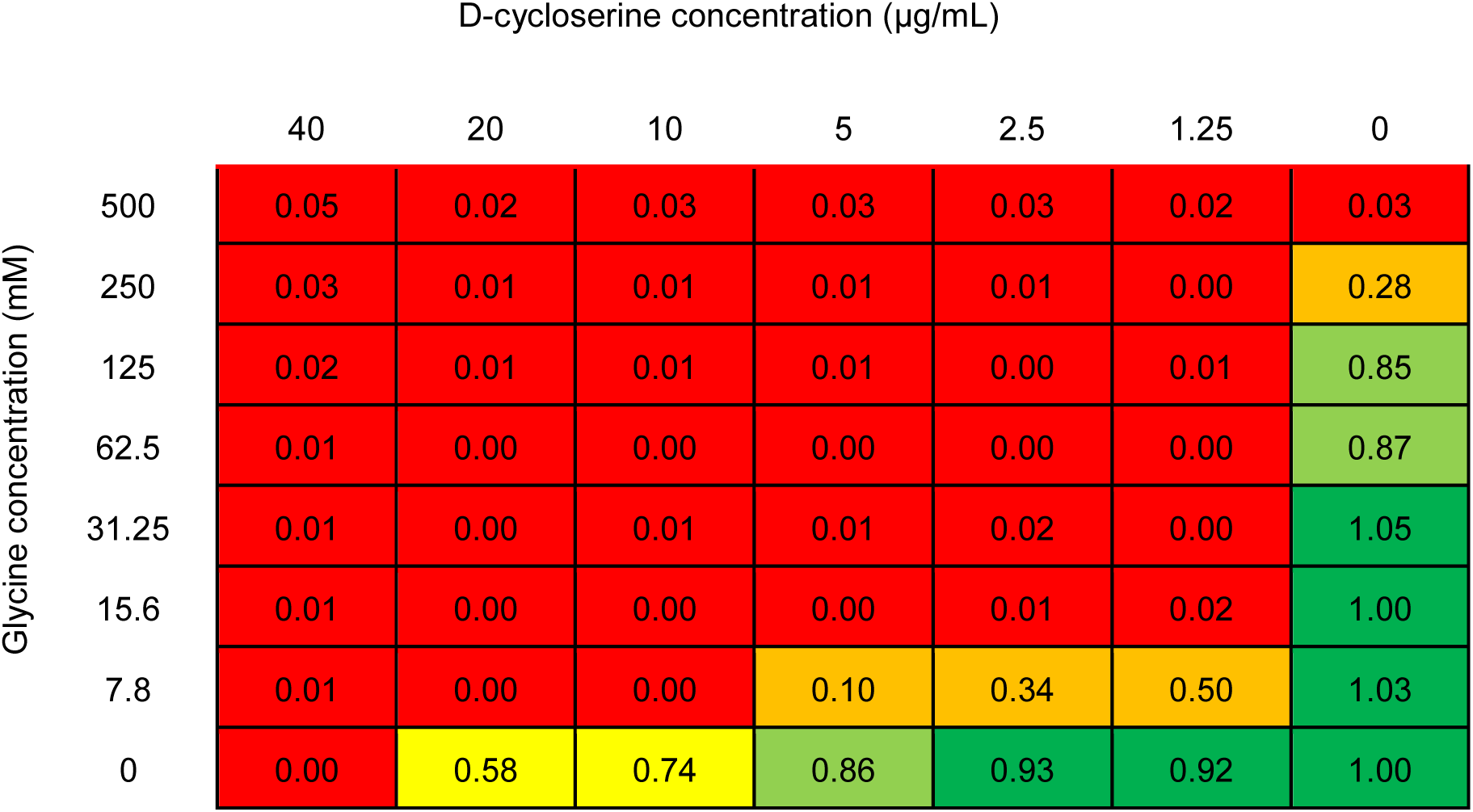
The alanine analog antibiotic D-cycloserine exhibits synergistic effects with excess glycine in inhibiting *S. aureus* growth. Growth of *S. aureus* strain JE2 after 15 hours in a checkerboard assay performed in TGM at 37°C with shaking. Results are reported as a fraction of the OD_600_ measured of the untreated control (bottom right, 0 glycine with 0 D-cycloserine). Wells in red exhibited total growth inhibition. Reported results are an average of three separate experiments.

### Glycine and D-cycloserine synergistically inhibit *S. aureus* growth

D-cycloserine (DCS) is an alanine analog antibiotic, and genetic inactivation of the alanine uptake transporter AapA results in hypersensitivity to DCS [46]. Thus, we predicted that if glycine inhibits alanine uptake at sub-lethal concentrations, combining glycine and DCS treatments should result in synergistic inhibition of *S. aureus*, even at relatively low glycine concentrations. To assess synergistic effects, a checkerboard assay was performed, combining D-cycloserine (MIC = 40 µg/mL) and glycine (MIC = 500 mM) treatments. In the checkerboard assay, the fractional inhibitory concentration index (FICI) was 0.09 of the combined treatments (<0.5 indicating synergism [48]), with inhibition of *S. aureus* in concentrations as low as 1.25 µg/mL DCS when combined with 15.6 mM glycine. Synergistic effects of the combined treatment were also observed in tryptic soy broth (FICI = 0.37), though the effect was somewhat diminished, likely due to high peptide and alanine content in this rich culture medium. A checkerboard assay with an *aapA* transposon mutant (*aapA*::tn), wherein the AapA alanine transporter is inactivated, showed hypersensitivity to DCS (MIC = 2.5 µg/mL), but no enhanced sensitivity to glycine (MIC = 500 mM) in individual treatments. However, DCS and glycine together were still synergistic against the *aapA* mutant (FICI = 0.26) (Figure S3). This result indicates that the mechanism of synergism observed between DCS and glycine is not limited to preventing uptake of alanine via AapA.

## DISCUSSION

In this work, we analyzed the genes necessary for *S. aureus* to withstand excess amounts of L-serine, glycine or diglycine. Our findings indicate that direct catabolism does not play a central role in coping with excess amounts of these amino acids; instead, resistance depends largely on maintaining uptake and synthesis of balancing amino acids, i.e., amino acids whose homestasis is disrupted by these treatments. Peptide uptake appears to play a key role in maintaining intracellular concentrations of these balancing amino acids upon glycine or L-serine treatment; in contrast, in the diglycine treatment, the only transporter that was important for growth was the free alanine transporter AapA. Alanine functions as the principal balancing amino acid upon glycine treatment, and the need for alanine uptake in diglycine was not surprising based on the mechanism of glycine toxicity. However, it was unclear why AapA is not important in excess glycine. We determined that this is because excess glycine interferes with alanine uptake, while peptide uptake appears unaffected. This is evidenced in part by the requirement for cells in excess glycine to convert D-alanine to L-alanine (via Alr), despite being provided with abundant extracellular L-alanine. This racemization becomes unnecessary if L-alanine-containing peptides are available to be taken up by DtpT. Direct measurement of alanine in culture supernatants further supports inhibition of alanine uptake by excess glycine. Wild-type cells treated with 1/5 the MIC of glycine consumed a significantly reduced amount of L-alanine from the culture medium. This newly discovered effect of excess glycine likely explains the necessity of alanine synthesis via Dat when glycine levels are high. Inhibition of alanine uptake by glycine also likely underlies at least in part the strong synergistic growth inhibition we observed with glycine combined with the alanine analog D-cycloserine (DCS). It should be borne in mind that DCS is not taken up by alanine transport systems, but competitively interferes with alanine metabolism in the cell [49, 50]. Genetic inactivation of alanine uptake by AapA has consequently been shown to cause hypersensitivity to DCS as a result of less intracellular alanine to offset toxic effects [46]. Thus, treatment with excess glycine phenocopies genetic inactivation of alanine uptake, rendering cells hypersensitive to DCS. However, we also found that DCS and glycine were synergistic in inhibiting an *aapA* mutant as well, hinting at additional mechanisms at play. It’s possible that this synergy persists in an *aapA* mutant because of glycine’s capacity to also prevent alanine uptake by alternative alanine transporters. However, it seems likely that much of the observed synergistic effect is because of the shared targeting of the D-ala-D-ala ligase (Ddl) enzyme by DCS and glycine; DCS antagonizes Ddl activity, while glycine is misincorporated in place of D-alanine by the same enzyme.

Other experimental results establish a strong connection between central metabolic activity and the capacity to resist excess amino acid toxicity. Our Tn-seq analysis indicates that disruption of the tricarboxylic acid (TCA) cycle and components of oxidative respiration confers a fitness advantage in all three amino acid treatment conditions. We suspect that the main benefit of inactivating the TCA cycle and respiration upon excess amino acid exposure is decreased oxidative stress, a phenomenon previously observed in the context of antibiotic tolerance and persistence [51, 52]. This would explain how these mutations protect against both excess L-serine and glycine, despite their distinct mechanisms of toxicity.

The involvement of central metabolism in amino acid toxicity is further supported by the need for the metabolic regulators encoded by *ptsH* and *ccpA* in excess diglycine or L-serine. Loss of *ptsH* (which encodes Hpr, a glucose phosphorelay component), or *ccpA* (which encodes a global glucose-responsive transcriptional regulator) results in defective glucose utilization, increasing dependence on alternative energy and carbon sources [36, 37]. It is tempting to assume that these genes are important for fitness in excess amino acids because their loss results in increased dependence on TCA cycle and respiration functions that we know are detrimental based on the discussion above. However, this model does not fully account for our observations, as the conditional essentiality of *ptsH* and *ccpA* is not seen upon glycine treatment. Therefore, we propose that loss of these genes results in selective induction of uptake systems that enhance influx of L-serine or diglycine, thus aggravating their toxic effects. It has previously been shown that expression of peptide uptake transporters such as DtpT are upregulated in response to nutrient starvation [37, 53]. Free glycine, however, may be constitutively taken up by *S. aureus,* which would explain why *ptsH* and *ccpA* mutants are not glycine-hypersensitive.

The idea that diglycine uptake—and not glycine uptake—is regulated, is consistent with several of our findings. First, the reduced sensitivity of wild-type *S. aureus* to diglycine (MIC > 1 M) compared to glycine (MIC = 500 mM) may be caused by repression of peptide uptake systems in nutrient-replete conditions, protecting from intracellular diglycine accumulation in early stages of growth when nutrient levels are high. Importantly, this is also when cell wall synthesis is at its peak and likely most sensitive to glycine misincorporation in peptidoglycan. Regulation of diglycine uptake by nutrient status is further evidenced by the phenotype of the *dat* mutant in excess diglycine, where sensitivity was only apparent in late exponential/early stationary phase of growth (Figure 4), at which point cell viability strikingly declined. In contrast, sensitivity to glycine results in a failure to initiate growth (Figure 3A). Taken together, the phenotypes of *S. aureus* in the presence of diglycine support the hypothesis that accumulation of diglycine is prevented by repression of uptake systems when nutrients are plentiful.

A notable result unique to L-serine treatment conditions is the essentiality of the TcyABC cystine/cysteine uptake system. The potential of excess L-serine to antagonize L-cysteine homeostasis is unsurprising given the structural similarities between L-serine and L-cysteine, but such a connection has not been previously reported. One explanation for this observation is that excess L-serine interferes with L-cysteine synthesis. Overwhelming L-serine concentrations (or O-acetylserine, given the deleterious effect on fitness of the serine acetyltransferase CysE) may act together as low-affinity L-cysteine analogs to inhibit the activity of enzymes involved in L-cysteine synthesis, some of which are normally subject to negative feedback inhibition by L-cysteine [39, 54]. It is unclear whether peptide uptake by DtpT (which was also critical upon L-serine treatment) is also important for supplying L-cysteine, or if the importance of peptide uptake is in supplying other balancing amino acids.

This work identifies new opportunities for rationally designed combination therapies. Our findings with combined glycine and DCS treatment illustrate the promise of such targeted approaches. Reducing the effective dose for DCS with a glycine adjuvant has therapeutic importance, as DCS side effects currently limit broader clinical use (reviewed in [55]). Likewise, therapeutic treatment with glycine alone has some promise, but its effective dose can be dramatically decreased by augmenting the treatment with DCS. Other novel treatment combinations informed by this study could pair glycine treatment with drugs that inhibit alanine synthesis, or pair L-serine with compounds that block cysteine uptake. Although a treatment to inhibit cysteine uptake has not been developed, targeting cysteine metabolism in bacteria is an area of active investigation [56, 57]. In considering other options for glycine combination treatments, ß-chloro-D-alanine disrupts alanine metabolism in a manner similar to D-cycloserine [58]. Its combination with glycine could enhance its therapeutic potential [59]. Other combination treatments may be enabled by our observation that glycine prevents alanine uptake.

Previous work has shown that genetic inactivation of alanine uptake results in hypersensitivity to cell wall-targeting antibiotics, including ß-lactams [46, 60]. Previous work has also shown that treatment with excess glycine sensitizes cells to cell wall-targeting antibiotics [13, 14]. Our finding reported here, that excess glycine inhibits alanine uptake, unifies these two previous observations and opens up new opportunities for the rational design of amino acid-based combination therapies.

## DATA AVAILABILITY

All data are available in this publication.

## ACKNOWLEDGMENTS

This work was funded by NIH grant RO1AI149491. We thank Tim Meredith for providing materials for creating *S. aureus* transposon mutant libraries. We thank Shawn Christensen for assistance in preparing and analyzing samples for metabolomic analysis.

